# Molecular basis of competence for neural induction in the chick embryo

**DOI:** 10.64898/2026.06.03.729870

**Authors:** Charlotte Colle, Vlad Arimia, Hui-Chun Lu, Claire Anderson, Leslie Dale, Claudio D. Stern

## Abstract

Competence is the capacity of a cell or tissue to respond to a specific inducing signal from a neighbouring tissue, by changing its fate in a specific direction. Neural induction is the process by which the epiblast of the early embryo responds to signals from the organizer (the tip of the primitive streak in amniotes) by forming a neural plate. Here we study why three regions of the early chick embryo lack competence to respond to neural induction by a grafted organizer: the outer anterior area opaca and the posterior area opaca at primitive streak stages (HH3^+^-4^-^), and the inner anterior area opaca at head process stage (HH5), in comparison with the competent anterior inner area opaca at HH3^+^-4^-^. Molecular analysis of these tissues, and their temporal dynamics following exposure to the organizer, reveals several differences. Among them, increased BMP and decreased ERK signalling characterise the non-competent regions. Inhibition of BMP can restore competence to HH5 epiblast; a combination of BMP-inhibition with ERK-stimulation by FGF8 can confer competence to the outer area opaca, whereas none of these can endow posterior epiblast with competence for neural induction. We conclude that spatiotemporal competence of epiblast for neural induction is regulated by several mechanisms, including extracellular signals.

## Introduction

In Developmental Biology, “competence”^1^ refers to the ability of a cell or tissue to respond to a particular inducing stimulus arising from neighbouring cells by adopting a particular fate. The concept was first introduced by Otto Mangold in the laboratory of Hans Spemann as *Reaktionsfähigkeit* (“ability to react”) (Mangold, 1932) and as *Reaktionsbereitsschaft* (“readiness to react”) by Machemer (1932) in the context of particular regions of the embryo, or of the same tissue at different times, that could respond or not to a graft of the “organizer” by forming an ectopic neural plate. The English word “competence” was introduced by Waddington (1932) and later elaborated further in a whole chapter in his book “Organisers and genes” (Waddington, 1940).

Waddington (1940) clearly explains that the concept of competence involves specificity of all of the following: the responding tissue (at a particular place and time during development), the inducing signal(s) (Waddington uses the term “evocator”) to which it responds, and the outcome of the interaction if the receiving tissue is competent, i.e. the result of the inductive interaction. He clarifies that for the concept to be valid, there must be at least two alternative outcomes, or fates, of the responding tissue.

Restricting the competence of cells to respond to particular inducing signals to specific places and stages is important because the number of possible cell fates vastly exceeds the small number of inducing signals, and even more so because many ligands share the same response pathways. Again it was Waddington who recognised this, noting that the extraembryonic region (area opaca) of the chick embryo is competent to respond to neural inducing signals from a grafted organiser (the tip of the primitive streak), and also reasoned that this implies that competence does not require the ectoderm to have been exposed to “endoderm” (hypoblast) (Waddington, 1934). He speculates that cells may have “successive competences”, for example first to form mesoderm, and then to give rise to neural structures, but that in some cases a cell might be competent to embark on different developmental trajectories when exposed to different inducers (Waddington, 1934; Waddington, 1940; Stern, 2000).

In his book “Epigenetics of birds”, Waddington states *“… little progress has been made with the embryological study of the waxing and waning of competence and the factors which bring it into being …”* (Waddington, 1956, p. 220). This statement is still largely true almost 70 years later, as we still know very little about the mechanisms that establish, or terminate, competence of the responding tissue for any inductive event during development.

Here we use Waddington’s original assay for neural induction in the area opaca of the chick embryo (Waddington, 1932; Storey et al., 1992; Stern, 2024) and take advantage of a recently defined Gene Regulatory Network for the responses of this ectoderm to neural inducing signals (Trevers et al., 2023). We analysed a competent region, the inner third of the anterior area opaca at Hamburger and Hamilton (HH) stage HH3^+^-4^-^ (Hamburger and Hamilton, 1951), which can respond to a grafted organizer (the tip of the primitive streak at HH3^+^-4^-^) by generating a neural plate that expresses *Sox1*, and compared it with three non-competent tissues: two in other regions of the area opaca at the same developmental stage but which are not competent at any stage during gastrulation, and the inner anterior region at a later stage (HH5), when this tissue has lost its responsiveness. We find that the lack of competence of two of these regions is due to differences in the levels of signals they receive: both have hyper-activation of BMP signalling, and one of them in addition displays low levels of MAPK/ERK activity. Adjusting the levels of these signals (inhibition of BMP alone in one case, and BMP-inhibition together with MAPK/ERK activation by FGF8 in the other) can confer them with competence. In contrast, the posterior part of the area opaca cannot be rendered competent by manipulating these pathways. We conclude that spatial and temporal competence in the embryo can be modulated by a variety of different mechanisms.

## Materials and Methods

### Neural induction assay

Fertile Bovan Brown Gold hens’ eggs (Henry Stewart, UK) were incubated at 38.5°C for either 16h, for the embryo to reach HH3^+^-4^-^, or 24h for HH5. Fine-grained staging was done following the criteria outlined by Selleck and Stern (1991). The embryo serving as host was then cultured ex ovo in modified New culture (New, 1955; Stern and Ireland, 1981). The tip of the primitive streak (the organizer) was excised from HH3^+^-4^-^ donors submerged in Tyrode’s saline solution (Tyrode, 1910), using 30G syringe needles. The dissected organizer was transferred to the desired location in the host using a P20 micropipette and grafted with its ventral (endodermal) side in contact with the host embryo, to avoid a contribution of the grafted node epiblast to the induced neural plate, as previously described (Storey et al., 1992; Streit and Stern, 2008; Trevers et al., 2023). When labelling of the graft was required, this was done by placing the excised organizer in a mixture of 1µl of 1mM DiI-Cell Tracker (Invitrogen C7001) in ethanol and 9µl of 60% (w/v) sucrose for 5 min. The dissected tissue was then rinsed in a drop of Tyrode’s saline before grafting.

Chick embryos at these stages of development are exempt from the requirement of a Home Office licence in the UK.

### Bead grafting

Heparin acrylic beads (Sigma H5263) were rinsed 3 times in PBS before being placed in either 25μg/ml in 0.1% BSA/PBS FGF8b (R&D Systems 423-F8-025/CF) or 0.1% BSA/PBS overnight at 4°C. After rinsing in Tyrode’s saline solution they were transferred to the desired location in the host embryo with a P10 micropipette and 30G syringe needle (Streit and Stern, 1999a; Streit et al., 2000).

### BMP inhibition: whole-embryo treatments

Embryos were placed for 5 min in 20μM DMH1 (Tocris 4126) in PBS/0.2% DMSO, or 0.2% DMSO in PBS, rinsed with Tyrode’s saline solution and incubated in modified New culture (see above) with either 2μM DMH1 in 0.02% DMSO, or 0.02% DMSO as control, in the albumen.

### Tissue collection and processing for RNAseq and NanoString nCounter

Dissected HH3^+^-4^-^ organizers were grafted to the desired location of the area opaca of a HH3^+^-4^-^ or HH5 host embryo for 5h (for RNA sequencing) or for 1h, 3h, 5h or 9h (for analysis by NanoString nCounter). Similar regions of the area opaca were also dissected from HH3^+^-4^-^ or HH5 embryos that had not received a graft (0h). Tissues were collected and processed as previously described (Trevers et al., 2018; Trevers et al., 2023; Lu et al., 2025). Twenty tissue pieces for each condition were collected for each sample, and samples were collected in triplicate. mRNA sequencing was done by the UCL Genomics Facility. For the 5h time point, organizer grafts that were in contact with competent or non-competent area opaca (triplicates of 20 organizers for each condition) were removed and processed in the same way for RNA sequencing. Prior to processing, tissue samples were stored at 4°C in RNALater (Invitrogen AM7020). For NanoString nCounter analysis, the area opaca tissue was dissected in the same way after either 1h, 3h, 5h or 9h incubation. For “uninduced” conditions, 4 pieces of tissue were collected for each biological replicate (experiments were done in triplicate for all regions), while for the conditions where the tissues were exposed to a graft, 6 pieces were collected in triplicate. Samples were collected and processed as previously described (Trevers et al., 2018; Trevers et al., 2023; Lu et al., 2025). For the generation of the heat maps from RNAseq data, the lists of ligands and receptors were generated based on the annotations in CellChatDB (Jin et al., 2021) and for transcription factors the annotation was based on the Human Protein Atlas (Uhlen et al., 2015) (https://www.v25.proteinatlas.org).

### In situ hybridization and immunostaining

In situ hybridization was performed as previously described (Streit and Stern, 2001). Immunostaining for pSMAD1/5/8 was performed using a protocol kindly provided by Sophie Brumm (Kruger et al., 2024). Embryos at stages HH3^+^-4^-^and HH5 were fixed in 4% paraformaldehyde in PBS for 1h at room temperature and dehydrated successively with 50% and 70% methanol in PBS/0.1% Triton-X11 followed by absolute methanol and stored at -20°C for up to 48h (Kruger et al., 2024). Samples were then rehydrated by the reverse process and placed into acetone at -20°C for 20 minutes. Embryos were then further permeabilised by incubating in 0.5% Triton-X100 in PBS for 4h at room temperature followed by blocking in 10% goat serum + 0.1%Triton-X100/PBS (“blocking buffer”) for 2h at room temperature. Samples were then incubated in 1:5000 pSMAD1/5/8 (Cell Signaling Technology 13820) or pERK1/2 (Cell Signaling Technology 9101S) primary antibody in blocking buffer overnight at 4°C on a rocking platform. They were subsequently washed with 0.1% Triton-PBS and incubated for 2h in 1:250 AlexaFluor 488-conjugated goat anti-rabbit IgG (Invitrogen A-11008) or AlexaFluor 456-conjugated goat anti-rabbit IgG (Invitrogen A-11011) secondary antibody in blocking buffer for 2h at room temperature with rocking. Then, embryos were washed with 0.1% Triton-PBS. DAPI staining for nuclei was performed by incubating the embryos for 10 min in 1:5000 DAPI in PBS at room temperature.

The staining intensity for pSMAD1/5/8 was measured using Fiji ImageJ (v. 1.54p) using the following pipeline: maximum intensity Z-projections were generated for both the DAPI and signal channels; based on the DAPI maximum intensity projection a binary mask was generated to contain only the nuclei. That selection was used on the signal average intensity projection to measure the fluorescence intensity in each particle with an area greater than 10μm^2^ . The mean fluorescence intensity values for each particle were then analysed. pSMAD levels were analysed from 6 embryos at stage 4^-^ for competent, outer and posterior regions, and 6 embryos at stage 5 for “late” epiblast, each analysed on the left and on the right sides. Each of these regions was captured using a 40x objective in a Leica SP8 confocal microscope as 4 tiles, to generate a 2x2 matrix.

### Sectioning and histological processing

The protocols were based on those described previously (Stern, 1993; Streit and Stern, 2001). Fixed embryos were washed with PBS containing 0.1% Tween-20 and dehydrated with absolute methanol for 10 min, isopropanol for 5 min and tetrahydronaphthalene for 30 min. Molten Paraplast (Sigma P3558) was added to make a 1:1 mixture with the tetrahydronaphthalene and embryos infiltrated for 30 min at 60°C. The embryos were further infiltrated in Paraplast at 60°C for 30 min 3 times before being placed into plastic moulds and left to set overnight. The wax blocks were sectioned at 10μm. The sections were then mounted onto glass slides coated with glycerol:albumen, floated on distilled water and then air dried on a warm plate. Dewaxing was done in Histoclear (National Diagnostics HS-200). Slides were coverslipped using either 3:1 Canada Balsam:Histoclear or CitiFluor AF1 (Electron Microscopy Sciences 17970-100).

### Imaging

Whole embryos were imaged using an Olympus SZH10 Stereomicroscope and histological sections imaged using an Olympus VANOX-T microscope. Images were captured using a QImaging Retiga 2000R Fast 1394 camera and QCapture Pro software (v. 7.0.2.22) and saved as 24-bit .tif files. Fluorescence images were superimposed on the corresponding brightfield frame using Fiji ImageJ. Specimens for fluorescence immunohistochemistry were imaged using a Leica SP8 confocal microscope and images saved as .lif files.

## Results

### Mapping competence for neural induction in the chick area opaca

Previous studies reported that the inner third of the area opaca is competent to respond to an organizer graft at stages HH3^+^-4^-^ but not at later stages, and that neither the posterior nor the outermost third of the area opaca is competent (Gallera and Ivanov, 1964; Gallera, 1971b; Gallera, 1971c; Gallera, 1971a; Dias and Schoenwolf, 1990; Storey et al., 1992; Streit et al., 1997; Streit and Stern, 1997). However, these conclusions were based on a diversity of criteria to identify the induced neural structures including morphology and different markers that may be difficult to compare to each other. To re-examine this using a uniform readout, we mapped the competence for neural induction by a grafted organizer by assessing whether the tip of the primitive streak (HH3^+^-4^-^) grafted at various positions can induce an ectopic region of expression of the pan-neural plate markers *Sox2* and *Sox1* (Pevny et al., 1998; Uchikawa et al., 1999; Wood and Episkopou, 1999) in the responding tissue. The results (Supplementary Figure S1) largely confirm those of previous studies (Gallera and Ivanov, 1964; Gallera, 1971b; Gallera, 1971c; Gallera, 1971a; Dias and Schoenwolf, 1990; Storey et al., 1992; Streit et al., 1997; Streit and Stern, 1997). We therefore selected four regions for further analysis and comparison: **“competent”** cells (inner third of the anterior area opaca at HH 3 -4) and three non-competent populations. Two that are never competent to respond to such a graft, the **“outer”** cells, in the outermost anterior third of the area opaca, excluding the outermost edge cells that adhere to the vitelline membrane (see Lee et al., 2022; Lee et al., 2025), **“posterior”** cells, in the inner posterior third of the area opaca and one (**“late”** epiblast), corresponding to the inner third of the anterior area opaca at HH5, which by this time has lost its competence to express neural markers in response to a graft of the organizer (Figure 1 A,B).

**Figure 1.**
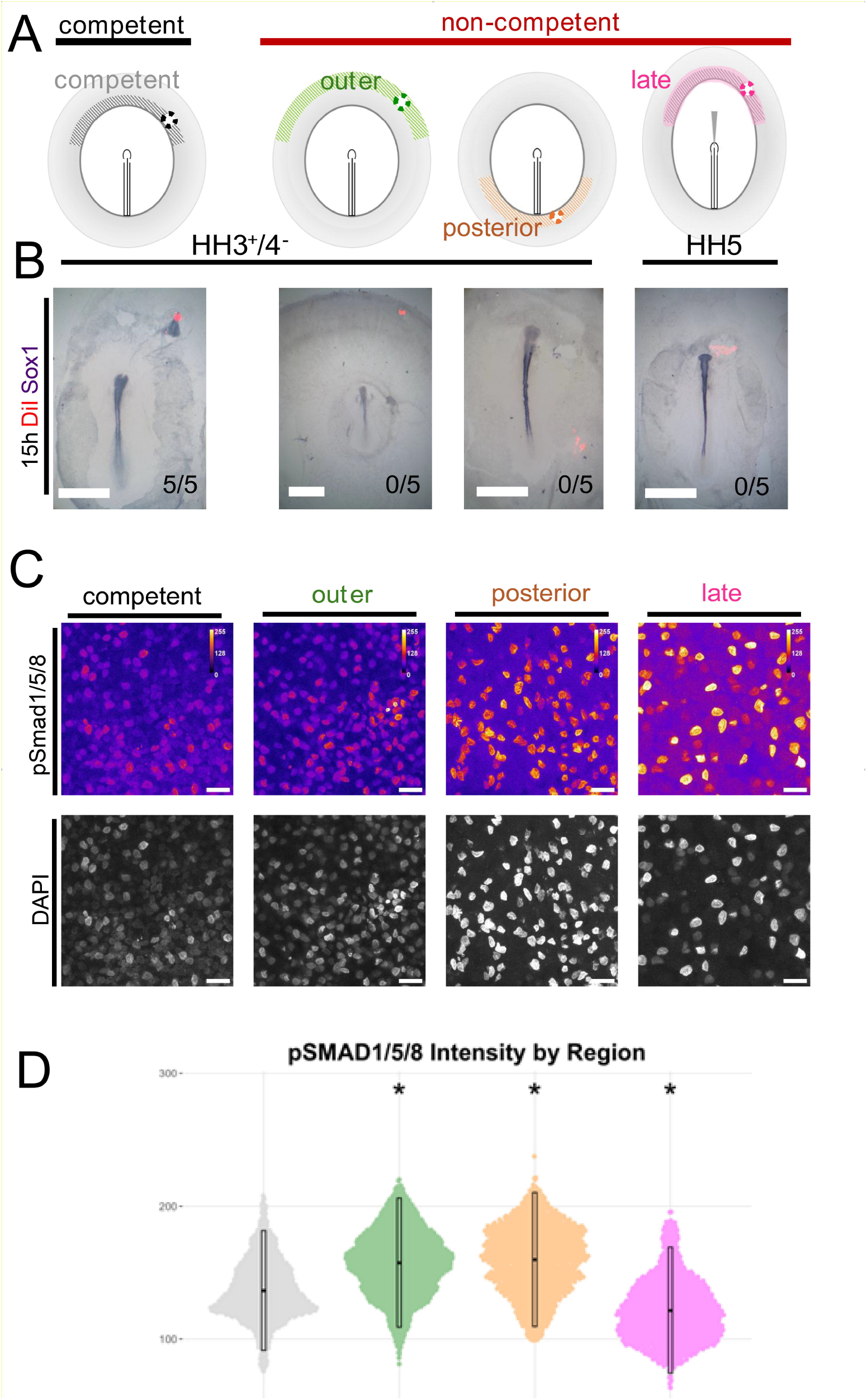
Regional differences in BMP signalling activity associated with competence or lack thereof. A DiI-labelled Hensen’s node was grafted onto different positions of the area opaca of primitive streak-stage embryos (HH3^+^–5) (A). After 15h, *Sox1* expression was assessed by in situ hybridization (B). *Sox1* expression was only observed in the first (“competent region”), the inner-anterior third of the area opaca at HH3^+^/4^-^ (5/5) (grey in A; representative induction in B). The peripheral “outer region” (green in A; induction assessed in B), and the “posterior region” (orange; induction in B) were not induced to express *Sox1* after 15h (0/5 for each in B). At stage HH5 (head-process stage), the inner-anterior third of the area opaca (pink; “late” cells) no longer responds to a HH3^+^-4^-^ node graft (0/5 in B). In each region, the phosphorylation levels of SMAD1/5/8 were assessed by immunohistochemistry, with representative images shown in C D and corresponding nuclear intensity quantification in D F. *: p<0.001 (Welch’s t-tests between competent and each non-competent region *: p<0.001). Scale bars: B, 500μm; C, 25μm.

To glean an initial view of molecular differences between the competent and the three non-competent tissues, we performed bulk RNA sequencing on the four populations. Comparison of the expression levels of transcription factors, receptors and members of signal transduction cascades (Supplementary Figure S2; Supplementary Data File 1) revealed that not only do the non-competent regions have a different transcriptional profile compared to the competent cells, but they also widely differ from one another. KEGG Pathway enrichment analysis (Supplementary Figure S3) of the regions indicated enrichment in some components and targets of ERK signalling in the competent cells, whereas mRNAs encoding components and targets of the BMP pathway appear to be more highly represented in the non-competent regions (Supplementary Figure S4). To test whether these differences, do indeed reflect differences in BMP activity between these tissues, we performed immunostaining against pSMAD1/5/8 (marking BMP signalling activity) (Figure 1C). Quantification of the staining intensity confirmed that BMP activity is higher in the outer and posterior cells than in the competent area opaca (Figure 1D).

### Many genes fail respond to an organizer graft in the non-competent positions

Next, we assessed the differences in the responses of the target epiblast to a graft of the organizer, in time-course (0h, 1h, 3h, 5h or 9h after the graft) between the competent region and each of the non-competent regions. We used a custom-designed NanoString nCounter probe set for 383 transcripts, previously used to generate a Gene Regulatory Network (GRN) for neural induction by an organizer graft (Trevers et al., 2023). Again, the three non-competent regions behaved differently to the competent cells and also to one another, with some responses diverging as early as 1h of exposure to the organizer, and increasing progressively thereafter (Supplementary Figures S5 and S6). After 9h many genes that are induced by an organizer graft in competent cells fail to be upregulated or repressed in the non-competent positions (Figure 2; Supplementary Data File 2) with a different subset of genes failing in each population. Collectively, our data show a gradual divergence between each non-competent tissue and the control, competent epiblast. PCA analysis of these transcription factors reveals the posterior cells to be most similar to the competent cells, followed by outer cells, while late cells have the most divergent transcriptional responses compared to competent ones (Supplementary Figure S6).

**Figure 2.**
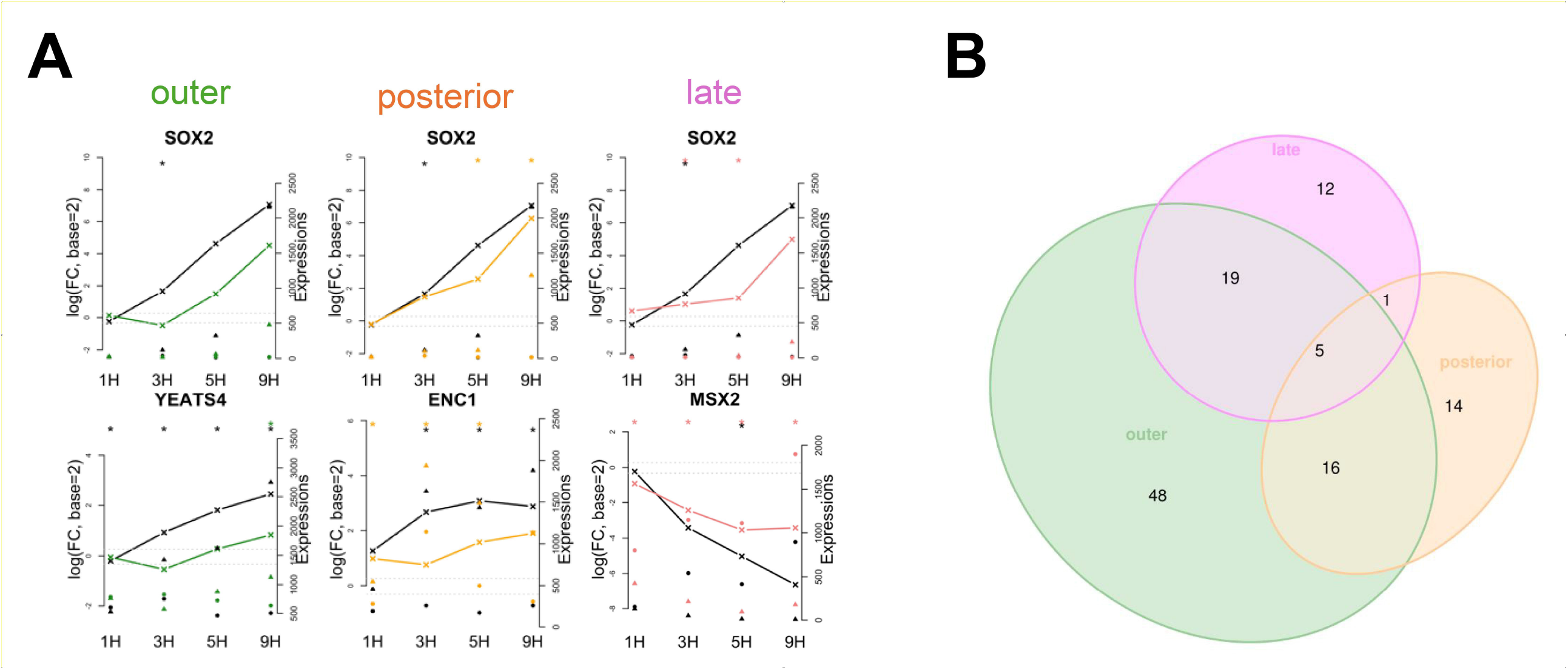
Some genes fail to be regulated appropriately in response to an organizer graft in non-competent regions. A. Time-course of expression (by NanoString; the y-axis (left) represents the log-fold-change relative to uninduced tissue for each region), and (right) expression level in FPKM of selected transcripts at 1, 3, 5 and 9h after a node graft, compared between the competent (black trace) and each non-competent (coloured) region. Experiments were performed in triplicate. Circles: FPKM mean values of uninduced tissues; triangles: FPKM mean values of induced tissues. *Sox2* (top row, all 3 regions), *YEATS4* (outer region; green) and *ENC1* (posterior; orange), are induced less strongly in the non-competent region in response to an organizer graft and *MSX2* (late region, pink) is an example of a gene whose expression fails to be downregulated by an organizer graft. The Venn diagram (B) summarises the number of response genes, from a total of 140 components of the GRN (Trevers et al., 2023), that behaves this way (“fail” to be up- or downregulated by the graft) for each region for at least one time point.

The datasets arising from the analyses in these two sections, were used find patterns that might reflect differences between competent and non-competent tissues, as revealed by different indicators. These can include pathway components or known targets, for example, so the specific results from each of these datasets cannot be compared with each other simply by looking at the names of the genes and the expression level. Thus, in some cases, different levels of a signalling activity are revealed by different levels of RNAs encoding agonists and/or antagonists, but in other cases we see different levels of target gene expression but not of the mRNAs for the players themselves, suggesting that in some instances the regulation of the pathway could be a post-transcriptional event. For these reasons, we only used them to point to possible causative signalling or other events that are amenable to direct experimental exploration, presented in the following sections of this paper.

### Inhibition of BMP activity rescues the lack of competence of “late” (HH5) cells

As mentioned above, RNAseq (Supplementary Figure S4) suggested a correlation between the loss of neural competence in late (HH5) inner anterior area opaca with elevated BMP activity, supported by pSmad1/5/8 immunostaining (Figure 1 C, D). We therefore investigated whether lowering BMP levels can rescue the loss of neural competence in HH5 ectoderm, by analysing the responses of the late region that had been treated with the Alk2/3/6 inhibitor Dorsomorphin-Homologue-1 (DMH1) (Hao et al., 2010) to an organizer graft, and assessing *Sox1* expression after 15 h. *Sox1* expression was not observed in DMSO-treated controls (0/6), whereas DMH1 treatment rescued the competence in 62.5% (5/8) cases, and some (3/5) of the induced structures had adopted a neural tube-like morphology (Figure 3). Histological sections confirmed that the ectopic *Sox1* expression was only in host epiblast adjacent to the graft. We also checked Smad1/5/8 phosphorylation levels after 15 h of treatment; in this case we do not see a significant decrease in BMP activity compared to DMSO controls (Supplementary Figure S7), suggesting that the effects of DMH1 on host epiblast are transient. These results show that the competence for neural induction by an organizer graft can be restored to the cells of the late (HH5) epiblast of the area opaca by lowering BMP activity, even for a relatively short time.

**Figure 3.**
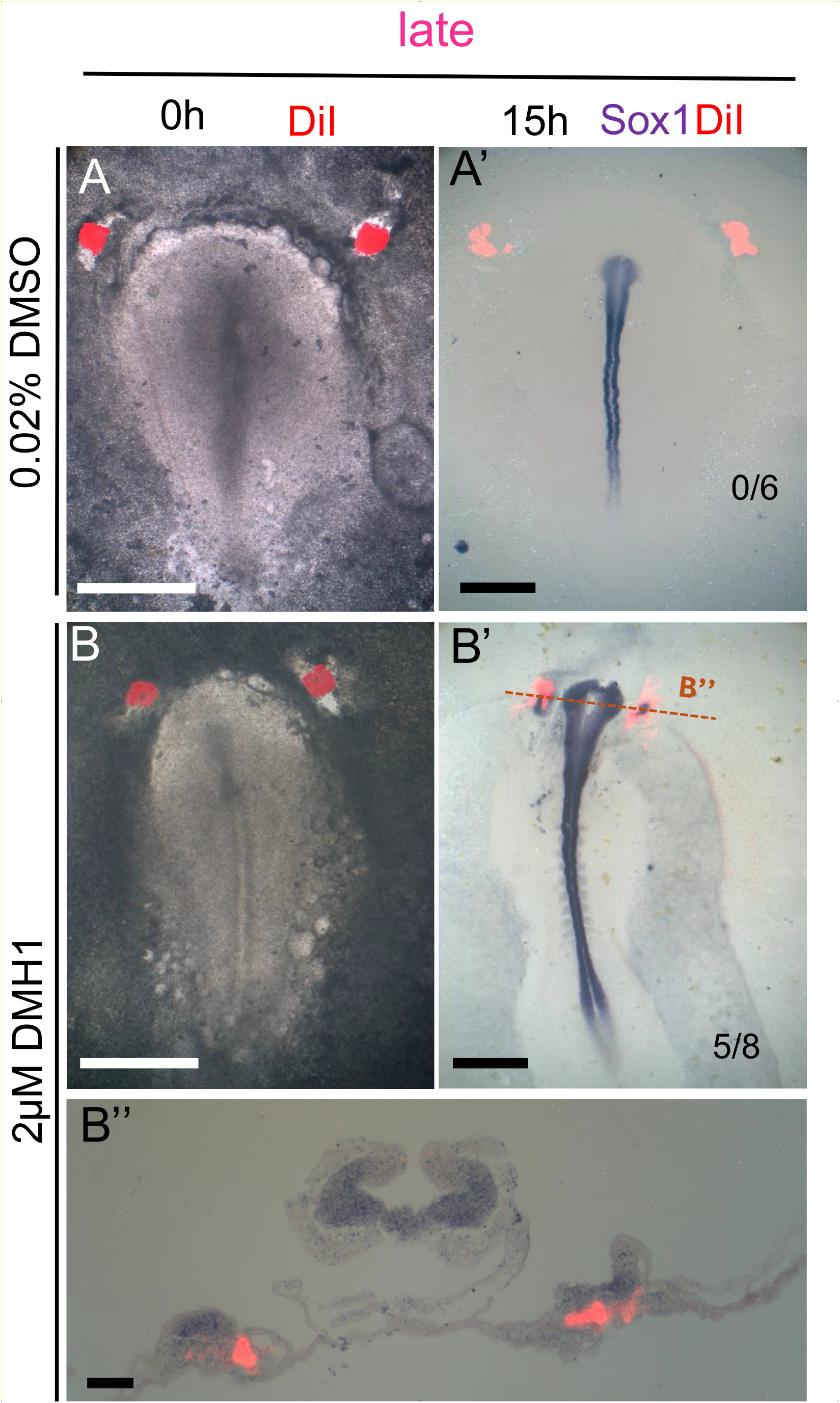
The loss of competence for neural induction at HH5 (“late” cells) is rescued by lowering BMP signalling activity. A DiI-labelled organizer graft was placed onto the inner area opaca of 0.02% DMSO (A) or 2μM DMH1 (B) treated embryos. No *Sox1* expression was observed near the grafts (0/5) in control embryos (A’), whereas DMH1 treatment allowed the graft to induce *Sox1* expression in epiblast adjacent to the graft (B’) (5/8). The induced structures also have a characteristic neural plate-like morphology as seen in transverse sections (B’’). Scale bars: 500μm for whole mounts, 50μm in B’’.

To explore this further, we attempted to inhibit BMP signalling in the responding tissue more directly, by electroporation of an expression construct encoding the inhibitory SMAD pathway component SMAD7 (de Almeida et al., 2008) before placing an organizer graft over the electroporated cells. When this was done in the “late” inner epiblast, competence to respond to the organizer graft was restored, with *Sox1* expression in 3/5 cases (Supplementary Fig. S8). This experiment confirms that manipulation of BMP activity in host cells can restore the loss of competence of late epiblast for neural induction, and also confirms that the rescuing effect of DMH1 treatment in the previous experiment is due to its action on the responding tissue rather than on the organizer graft.

### The “outer” cells can be endowed with competence by lowering BMP and exposure to FGF8, but not by either of these treatments alone

Unlike the late (HH5) area opaca epiblast, lowering BMP signalling activity by DMH1 treatment does not confer competence for neural induction to either outer or posterior cells (Supplementary Figure S8). Since both of these non-competent populations appear to have lower levels of MAPK/ERK signalling than the competent area opaca (Supplementary Figure S3), we tested whether stimulating ERK (a target of FGF, EGF and related ligands) could confer competence to these regions. To this end, we placed two FGF8-soaked heparin acrylic beads into the outer or posterior region, flanking an organizer graft. No *Sox1* induction was observed after 15 h (Supplementary Figure S10) in either case (0/10, 0/10, respectively). We also performed the same experiment on late hosts (HH5), and again there was no *Sox1* induction (0/6) (Supplementary Figure S10).

Next, we tested whether inhibition of BMP activity together with ERK1/2 stimulation can endow the outer region with competence for neural induction by a grafted organizer using DMH1 together with FGF8-soaked beads adjacent to an organizer graft; this led to *Sox1* induction by an organizer graft in outer area opaca epiblast (4/6) (Figure 4). Thus, the competence for neural induction by the organizer can be extended to the outer area opaca cells at stage HH3^+^-4^-^ by increasing ERK activity together with inhibition of BMP activity, but not by either treatment alone.

**Figure 4.**
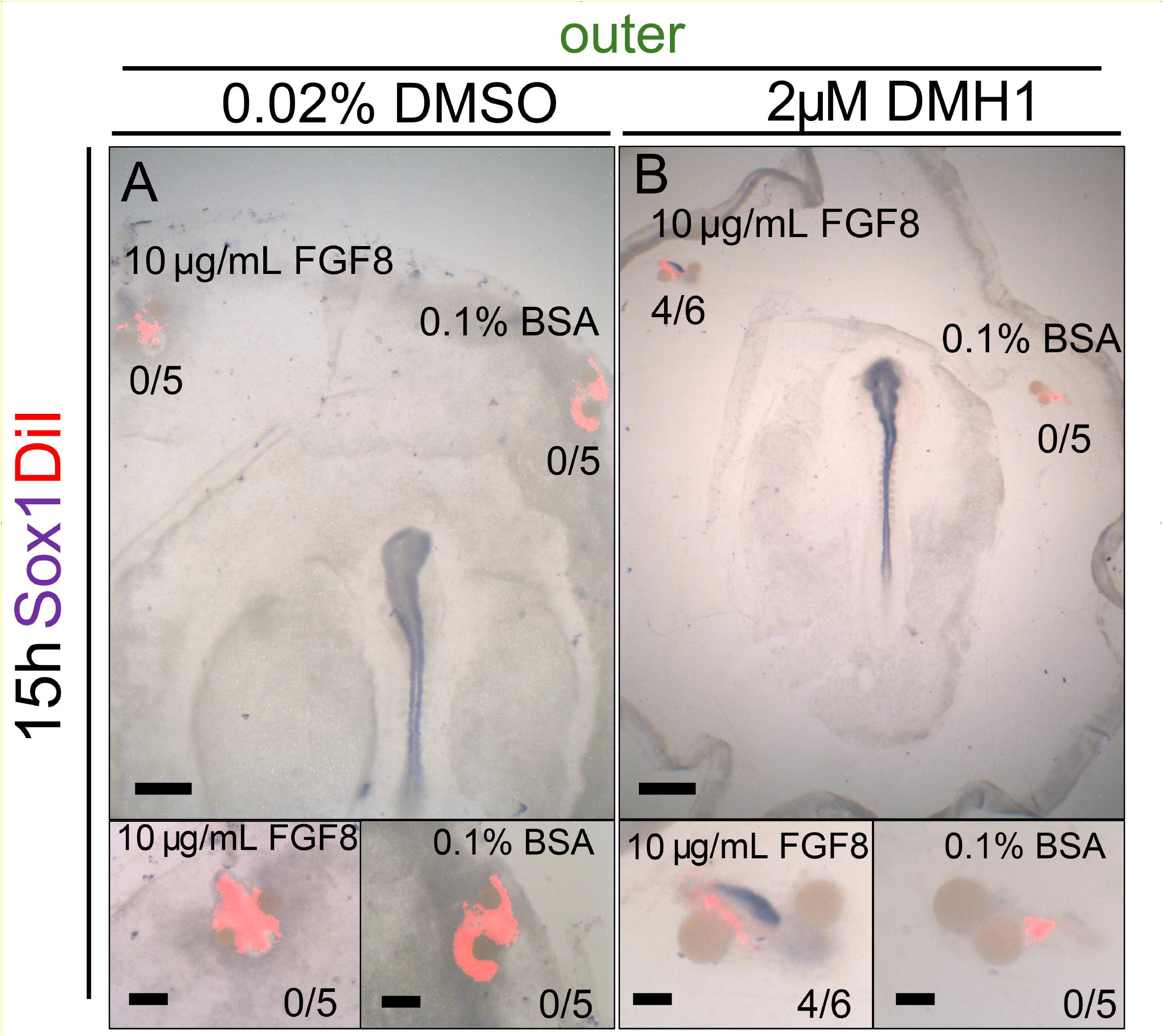
FGF treatment together with BMP inhibition confers competence to “outer” cells. A node was grafted onto the outer region of a host embryo treated with either control 0.02% DMSO (A) or 2μM DMH1 (B), flanked by beads soaked either in 10μg/ml FGF8 or 0.1% BSA control. In the 0.02% DMSO controls no detectable *Sox1* expression was observed adjacent to the grafts either in organizer+0.1% BSA beads (0/5) or for organizer+10μg/ml FGF8 beads (0/5). In 2μM DMH1-treated embryos, no *Sox1* expression was seen next to the organizer + vehicle control grafts (n= 0/5), but *Sox1* was induced when the organizer graft was flanked by 10μg/ml FGF8 beads (4/6). Scale bars: 500μm in A-B, 50μm for close-up views below.

### Neither BMP inhibition, nor FGF8, nor both treatments together, can confer neural competence to “posterior” cells

In contrast, neither DMH1 treatment alone (0/5) (Supplementary Figure S8) nor FGF8 (0/5) (Supplementary Figure S9), nor the combination (0/5) could rescue the competence of posterior area opaca epiblast (Figure 5).

**Figure 5.**
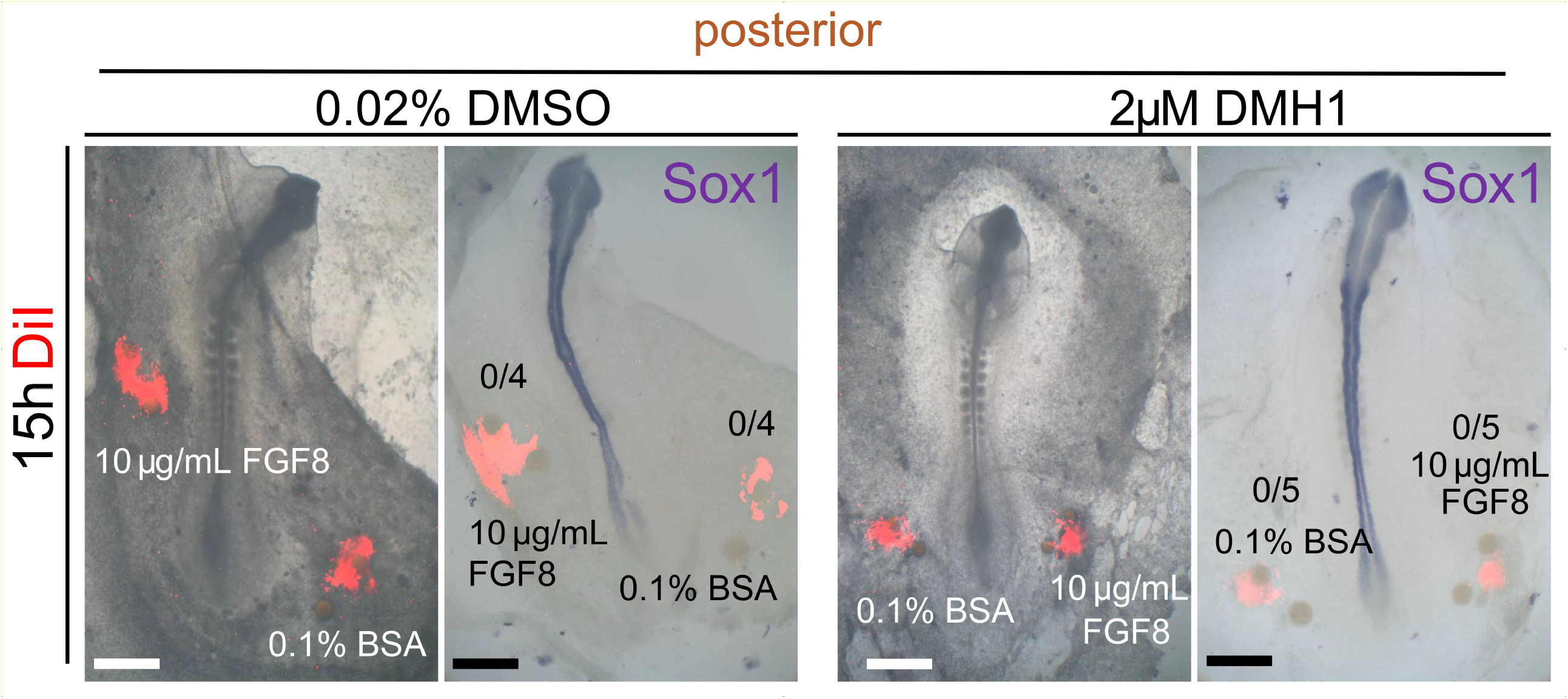
Co-treatment of BMP inhibition and FGF stimulation does not endow neural competence to the “posterior” cells. DiI-labelled organizers were grafted onto the posterior region, flanked by two heparin acrylic beads soaked either in control (0.1% BSA) solution or 10μg/ml FGF8. Following 15h of incubation either in control (0.02% DMSO) or BMP inhibitor treatment (2μM DMH1) the expression of *Sox1* was assessed by in situ hybridization. None of the tested conditions was able to confer neural competence to the posterior cells (DMSO controls 0/5 in combination with BSA soaked beads and 0/4 in the presence of FGF8 beads; DMH1 treatment with BSA beads 0/5 and 0/5 in combination with FGF8 beads). Scale bars are 500μm.

Taken together, these experiments show that different mechanisms can regulate competence for neural induction by the organizer in different non-competent regions: the late (HH5) inner area opaca epiblast can be rescued by lowering BMP, the outer area opaca epiblast can acquire competence by simultaneous inhibition of BMP and stimulation of ERK with FGF8 (but neither of these treatments alone), and the posterior area opaca cannot be rendered competent by any of these treatments, either alone or in combination.

### The node maintains its organizer properties upon grafting onto a non-competent territory

In addition to properties of the non-competent cell populations that could affect their responses to signals from the organizer, it is conceivable that the environment (for example through high levels of BMP at the graft site) could affect the organizer graft itself, causing it to lose its expression of critical signalling molecules (Stern, 2025). To test whether the organizer properties of the node are affected by exposure to non-competent territories of the area opaca, we designed a serial grafting experiment (Figure 6A). The organizer was initially grafted onto either the competent region, or one of the non-competent ones. After 5h of incubation, the node was removed and re-grafted onto a competent territory in a new host embryo. Grafted embryos were assessed for induction of the mature neural plate marker *Sox1* as before (Figure 6). *Sox1* expression was observed regardless of whether the node had initially been grafted onto competent (4/4), posterior (4/4), outer (3/4) or late (2/2) cells. Thus, the graft maintained its organizer properties regardless of the territory of the area opaca even if it had previously been exposed to a non-competent region of epiblast.

**Figure 6.**
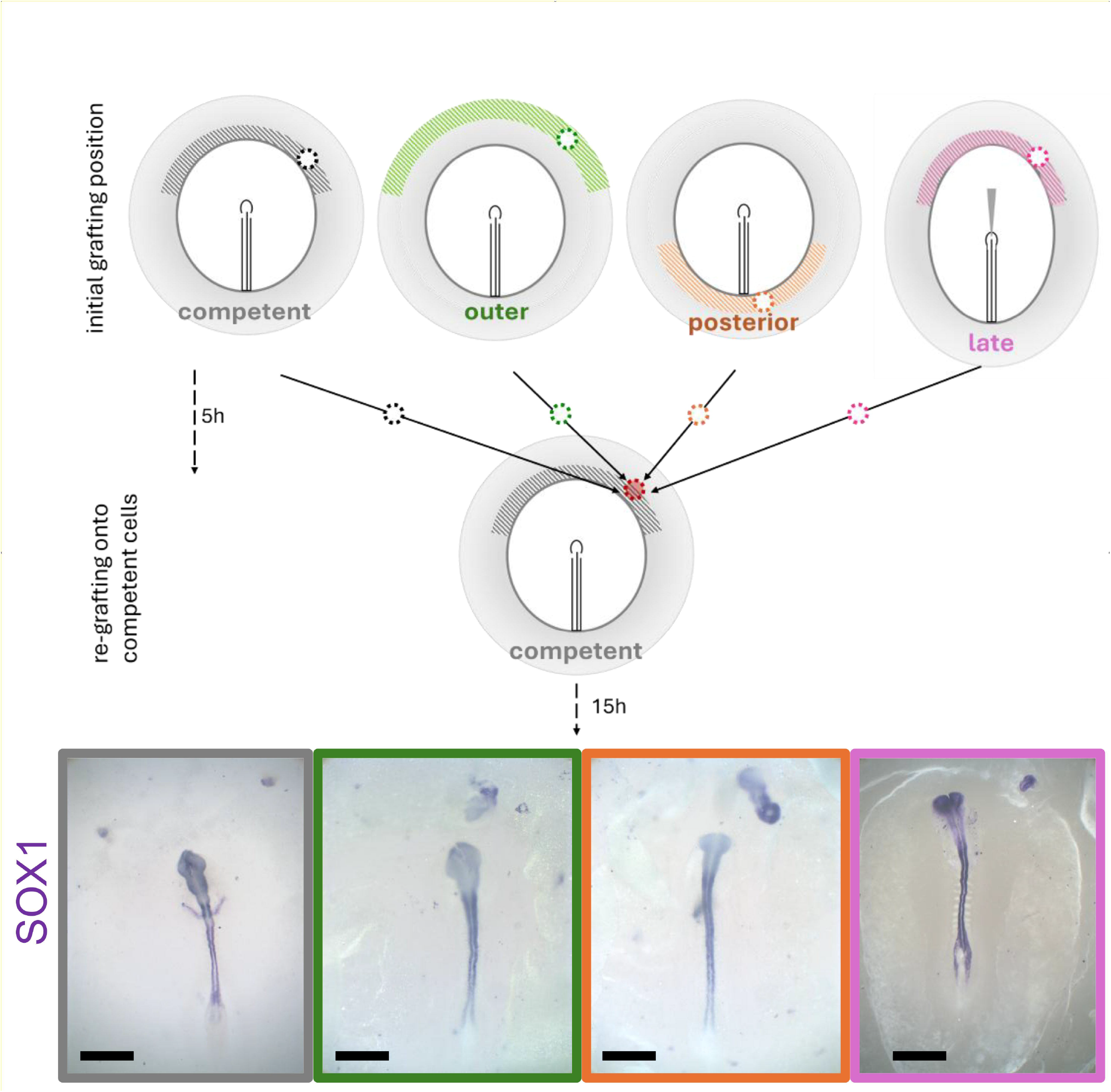
Lack of competence is not due to inhibition of the inducing ability of the organizer graft by the host environment. A serial grafting experiment was designed (A) to test whether exposure of the organizer to non-competent territories affects its ability to induce a neural plate. A node was grafted onto one of the four regions and removed after 5h, then grafted onto competent cells of another host embryo at HH3^+^-4^-^ . *Sox1* expression was observed whether the node had initially been grafted to competent cells (3/3), onto the posterior region (4/4), outer region (3/4) or late cells (3/3). Scale bars: 500μm for whole mounts, 100μm for sections. Each of the regions in which the graft had originally been placed is colour coded in the diagrams above, and this colour used to highlight the border of each of the cases in the results shown below: competent: grey; outer: green; posterior: orange; late: purple.

As an additional indication of whether the node itself is affected by the environment into which it is grafted, we performed RNAseq to compare the transcriptome of nodes that had been placed into each of the four regions, after 5 hours. We find that although there are some differences, the expression of node markers such as *GSC (Goosecoid), ADMP, GNOT2* and *NOTO* is maintained and the grafts still express signalling molecules such as *CHRD* (*Chordin*) and FGFs (Supplementary Figure S11). The only significant differences observed were a decrease of *ADMP* expression in organizers that have been exposed to the late cells and an increase in *GNOT2* mRNA for grafts placed onto the outer region (Supplementary Figure S11).

## Discussion

In principle, the responsiveness of a cell to a particular extracellular signal is dependent on its expression of appropriate receptors for that signal, of components of the intracellular response pathway, chromatin accessibility of targets of the pathway, and on its “state” as defined by the combination of factors (including transcription factors and other components) set up during its earlier history. Therefore, gain or loss of competence in particular tissues during development could be regulated by changes in any of these. In “Organisers and genes” (Waddington, 1940), Waddington speculates that loss of competence is “autonomous” to the tissue, but later he comments: *“It is … difficult to believe that all competences throughout development can arise autonomously, without dependence on previous inductive processes.”* (Waddington, 1956) (p. 220), realising the importance of the cell’s prior developmental history. The debate between autonomous and extrinsically regulated changes in competence has been the subject of debate ever since. In studies on lens induction in amphibians, Servetnick and Grainger (1991) argued for entirely autonomous, time-regulated processes, but other studies have implicated external signalling molecules like Hepatocyte Growth Factor (HGF) (Streit et al., 1997; Streit and Stern, 1997), and even changes in mechanical tension in the responding tissue due to hydrostatic pressure (Alasaadi et al., 2024) in the regulation of competence for other inductive interactions.

While it is possible to study the mechanisms by which a particular competence is lost over time in a given tissue, the time at which that competence is acquired by the tissue is impossible to determine experimentally. This is because when the organizer is grafted at an earlier stage (prior to about HH2-3), it induces mesoderm from the host epiblast; for example, see Stern et al. (1995). Although some neural tissue may form later, it is impossible to determine whether this is a separate neural induction or whether it is indirectly elicited by the induced mesoderm. In fact, there may not be a time at which competence is established, if the developmental progression simply involves either a progressive loss of potential to give rise to particular structures, or changes in the competence of the tissue over time such that the same cells will give rise to different tissues according to the time of exposure even to the same signal(s). Indeed, Waddington noted both of these features in several of his writings.

In the present study, the HGF receptor *MET* is expressed at extremely low levels in all tissues (Supplementary Data File 1 – 0h time point). However, this ligand activates several pathways downstream of the receptor, including ERK (shared with FGF, EGF, IGF and others) and NFκB (also involved in the regulation of BMP signalling (Lee et al., 2024)). Our study implicates both of these pathways in the lack of competence for neural induction in the outer area opaca (BMP+ERK) and in the late area opaca epiblast (BMP), although the endogenous triggers of these pathways that underlie the regulation of competence in this epiblast are unclear. In the case of BMP, however, it has been extensively documented that the expression of BMP4 itself is high in the posterior part of the embryo including the area opaca, and is also higher throughout the area opaca at HH5 than at primitive streak stages (HH3^+^-4^-^) (Streit et al., 1998; Streit and Stern, 1999b; Chapman et al., 2002; Linker and Stern, 2004).

A study of responses of chick and mouse ventral hindbrain neuroepithelial cells to TGFβ signals emanating from the neighbouring isthmus, which acts as the midbrain-hindbrain organizer (MHB), revealed that the competence of the responding cells is regulated by activation of canonical SMAD2/3, functioning as a temporal switch, terminating early motor neurone competence and enabling competence to give rise to serotonergic neurons (Dias et al., 2014b). This is reminiscent of Waddington’s “successive competences” (see above) and a further example that signalling through SMADs can modulate competence to inductive interactions.

We have not been able to identify a clear molecular basis behind the lack of competence of the posterior area opaca. This region also expresses high levels of BMP4 (Streit et al., 1998; Streit and Stern, 1999b; Dias et al., 2014a), but inhibition of this ligand does not restore competence. However, receptors for some ligands previously shown to be expressed in the organizer (Lu et al., 2025) such as the *Midkine* (*MDK*) receptor *PTPRζ1* (*PTPRZ1*), or the Somatostatin (SST) receptors *SSTR1/2/4/5*were observed to be downregulated in the posterior area opaca compared to the competent region (Supplementary Figure S2; Supplementary Data File 1) raising the possibility that these signalling pathways also play a role in regulating competence for neural induction. However, the lack of reagents makes it impossible at this stage to design clear functional experiments to test this. Changes in chromatin conformation or other factors affecting accessibility of targets in the neural induction process could also be involved, but these are not easily revealed by analysis of the transcriptome.

The recently described GRN depicts 5,614 interactions between 175 transcription factors whose expression either increases or decreases significantly in competent area opaca cells following exposure to a graft of the organizer, with fine temporal resolution (1-2 h) covering the period from 0-12 h after grafting (Trevers et al., 2023). When the responses to a grafted node into each non-competent position are compared to a graft into the competent area at the 3 h time point (see above and Figure 2, Supplementary Figure S6 and Supplementary Data File 2), we find that several genes fail to be up-or downregulated significantly by the graft in one or more non-competent locations as compared to the competent region (summarised in Supplementary Data File 3 and Supplementary Figure S12). After adjusting for expression levels, the degree of difference between competent and non-competent, and selecting those transcription factors that have direct targets in the GRN, two genes stand out that fail to be upregulated by a node graft at this time point: *BLIMP1* (*PRDM1*) fails to be upregulated in outer and posterior non-competent regions, and *MAFA*, which fails to be upregulated in late epiblast. Supplementary Figure S12 shows the GRN subnetworks downstream of each of these transcription factors. BLIMP1 can be an activator or a repressor. At the 3h time point, *BLIMP1* represses numerous genes that are expressed at 0 h, as well as SIX1, which is first expressed at 12 h. BLIMP1 positively regulates *ZIC3, OTX2, GBX2, SOX3,* and *TCF7L1*. MAFA can also act as activator or repressor. It represses many genes that are expressed at 0 h such as *CEBPB, IRF7, KLF6, and NFKB2*, as well as *SIX1*, which is first expressed at 12H; as an activator, it positively regulates many genes including the critical pan-neural plate marker *SOX2*. Therefore, BLIMP1 and MAFA could be responsible, alone or in combination with other variables, for the lack of competence in posterior and outer cells, and in late epiblast, respectively.

During very early development, *BLIMP1/PRDM1* is expressed broadly in the embryonic epiblast but not in the epiblast of the area opaca. Upon exposure to either a node or to other inducing tissues like head mesoderm (which can induce a pre-placodal state), competent area opaca epiblast starts to express this gene within 3 h in response to either tissue, along with other genes that have similar expression profiles in early development, including *ERNI, OTX2* and *SOX3* (Trevers et al., 2018). This initial response of the area opaca competent epiblast to inducing signals (“pre-neural/pre-border/pre-placodal“, or “common state”) has been likened to a return to a state similar to the pluripotent state of embryonic stem cells and early epiblast, before the induced tissue embarks on a new direction in response to the inducing stimulus (Trevers et al., 2018; Prajapati et al., 2019). At least two of the “common state” genes, *ERNI* and *PRDM1*, are first induced but then need to be downregulated for neural induction to proceed (Papanayotou et al., 2008; Prajapati et al., 2019). FGF signalling from the inducing tissues seems to be the main signal triggering this state (Streit et al., 2000; Trevers et al., 2018; Lu et al., 2025).

FGF signalling can phosphorylate the linker region of Smad1 and in this way contribute to inhibition of Smad1-dependent BMP signalling (Pera et al., 2003). However, the ability of FGF to regulate the early “common state” appears to be independent of its ability to inhibit BMP (Linker and Stern, 2004; Trevers et al., 2018; Lu et al., 2025). To what extent the acquisition of competence by outer area opaca by BMP-inhibition together with FGF is due to each of these two mechanisms is difficult to establish.

Clearly, the ability of cells to respond appropriately to an inducing signal and choosing one of two (or more) outcomes requires all of the conditions to be satisfied, therefore there may be multiple reasons for the initial acquisition and later loss of competence by different responding tissues. Waddington himself recognised the underlying complexity of inductive interactions as early as 1934: *“… it might be supposed that during the period of competence to form X, there is present in the competent tissue a certain constellation of chemical substances or processes which are in unstable equilibrium, so that the addition of small quantities of other substances (the organiser substances) can cause the whole equilibrium to break down, and the system to alter progressively in a certain direction towards some new equilibrium”* (Waddington, 1934). However, our study reveals that reducing the level of BMP signalling in one case (late inner area opaca epiblast) and this together with stimulating ERK signalling using FGF8 in another (the outer area opaca epiblast) is sufficient to confer competence to these two tissues, suggesting that changes in components of these signalling pathways could be the major mechanism accounting for their lack of responsiveness in a neural induction assay.

## Acknowledgements

This study was funded by grants from NIH (R01 MH60156) and BBSRC (BB/R003432/1 and BB/K007742/1) to CDS. CDS was a Wellcome Trust investigator (107055/Z/15/Z). CC was a student in the Wellcome Trust funded PhD programme “Developmental and Stem Cell Biology” at UCL. We are grateful to Dr Sophie Brumm for sharing her protocol for immunostaining of pSmad1/5/8.

## Data availability

all data generated by this study is included in the Supplementary Datasets. RNAseq data have been deposited in GenBank with accession number: XXXX

## Artificial Intelligence

We have not used any AI tools for any part of this paper, including figures, writing or editing.

## Author contributions

CC, VA and CA: performed experiments; H-CL: carried out all analysis including bioinformatics and statistical tests; LD: discussions, reviewed manuscript; CDS: project design, overall supervision, obtained funding. VA and CDS wrote the first version of the paper. All authors reviewed and approved the manuscript.

## Footnotes

1 Note: the correct word in the embryological and developmental biology literature is “competence” rather than “competency”. This is not only for historical reasons, as the word defined originally by Waddington, but also because the former implies a binary response – the responding tissue may or may not respond to the stimulus, whereas the word “competency” is more often used to describe an ability that can vary by degrees.

