## Supplementary Figures for "Molecular basis of competence for neural induction in the chick embryo"

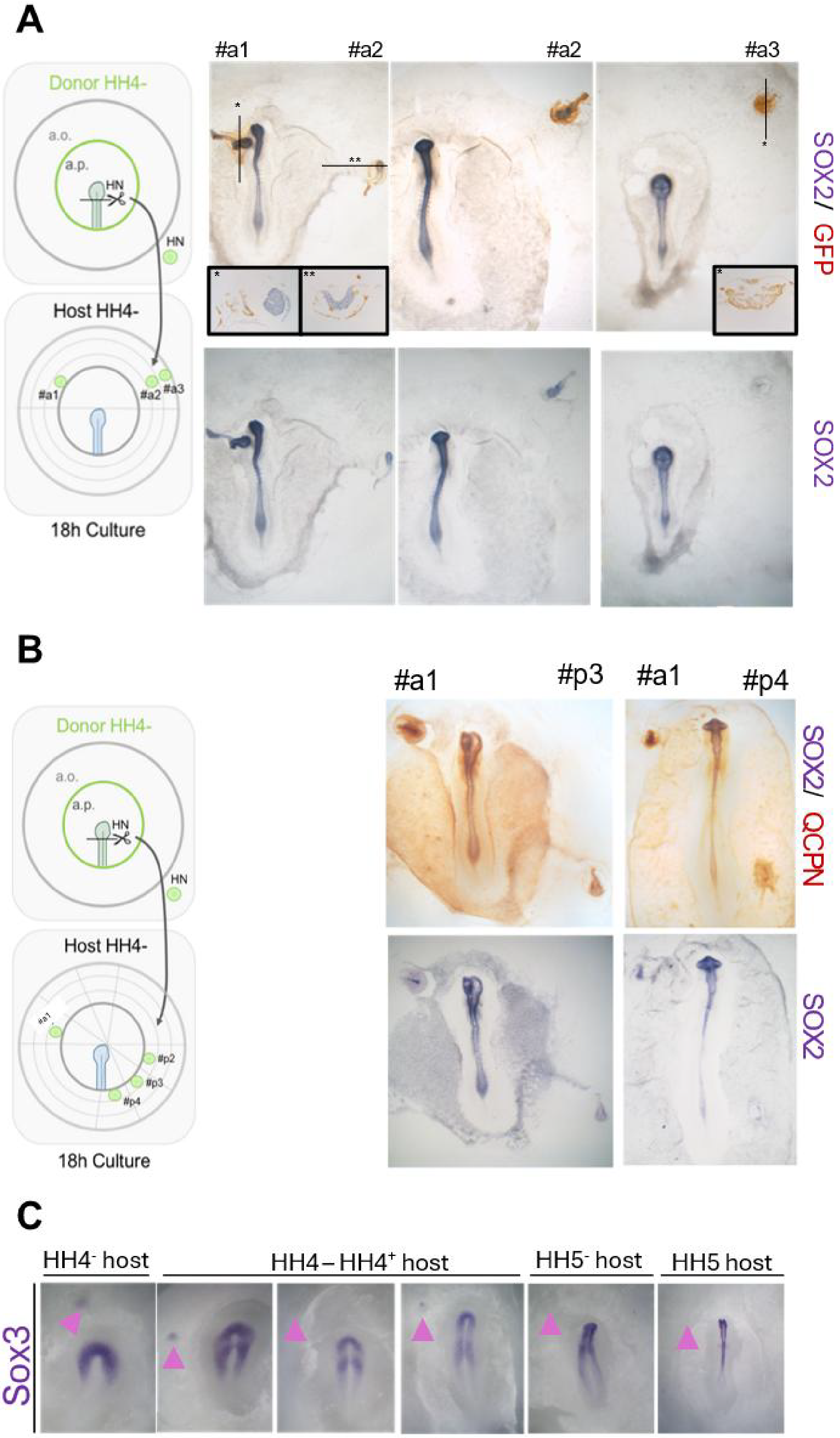
A
Donor HH4-
a.o.
a.p.
HN
HN
Host HH4-
#a1
#a2
#a3
18h Culture
#a1
#a2
#a2
#a3
SOX2/ GFP
SOX2
B
Donor HH4-
a.o.
a.p.
HN
HN
Host HH4-
#a1
#p2
#p3
#p4
18h Culture
#a1
#p3
#a1
#p4
SOX2/ QCPN
SOX2
C
HH4- host
HH4 – HH4+ host
HH5- host
HH5 host
Sox3

**Supplementary Figure S1 – Mapping competence for neural induction in the area opaca** Grafts of the amniote organizer, the anterior tip of the primitive streak of stage $HH3^{+}$-$4^{-}$ donor embryos (either from a GFP-transgenic donor or from a quail donor embryo), were placed onto different regions of the area opaca of wild-type chick host embryos to test the competence of the latter to neural induction. For this, in stage $HH3^{+}$-$4^{-}$ host embryos, the anterior area opaca (A) was tested by grafting onto the inner (#a1), middle (#a2) or outer (#a3) thirds at the level of the host Hensen's node, and the posterior area opaca (B) was assessed by grafts placed in the inner third of the area opaca either at the "4 o'clock" position below the level of the host node, (#p3) or at the level of the base of the streak (#p4). The distinction between the anterior and posterior area opaca was made based on the level of the host Hensen's node with an imaginary horizontal line passing through the node separating the two regions. After overnight incubation the embryos were assessed by in situ hybridization for *Sox2* (A, B) or *Sox1* (not shown) and the grafts identified using either anti-GFP immunostaining (A) or QCPN (anti-quail nucleolar antigen) immunostaining (B) as appropriate to the donor tissue used. Embryos were sectioned to assess the contributions of the node and host to the ectopic structures (* and **). To determine the temporal window of competence, a time course experiment was performed (C) where $HH3^{+}$-$4^{-}$ organizers were grafted onto the inner anterior third of the area opaca of hosts of various stages starting from $HH4^{-}$ to HH5. After overnight incubation the expression of early neural plate markers (*Sox1*, *Sox2* or *Sox3*) assessed by in situ hybridization. This example shows *Sox3* expression. Arrowheads indicate the graft sites. The organizer does not induce ectopic neural plate marker expression in hosts older than stage HH4. a.o. – area opaca; a.p. -area pellucida

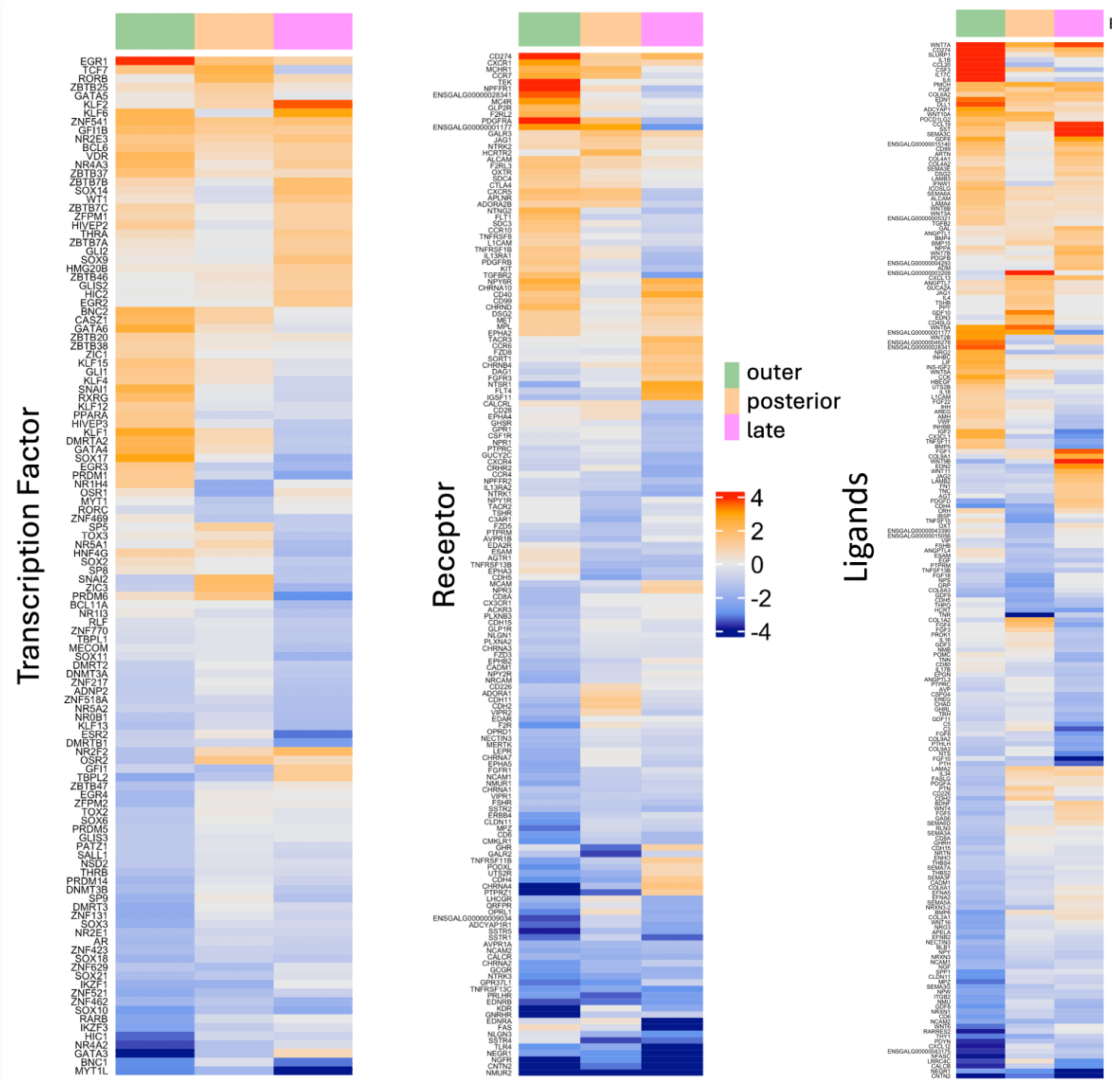


**Supplementary Figure S2 – Transcriptional profile of the non-competent regions.** Heat maps (left to right) show the z-score-scaled fold-change of non-competent vs competent regions for transcription factors, receptors, and ligands selected from the RNAseq comparisons. Genes were selected based on absolute $\log_2$(fold change) >1 in at least one non-competent region. Additionally, for upregulated genes, an fpkm threshold >10 was required in the relevant non-competent region, and for downregulated genes, fpkm>10 was required in the competent region.

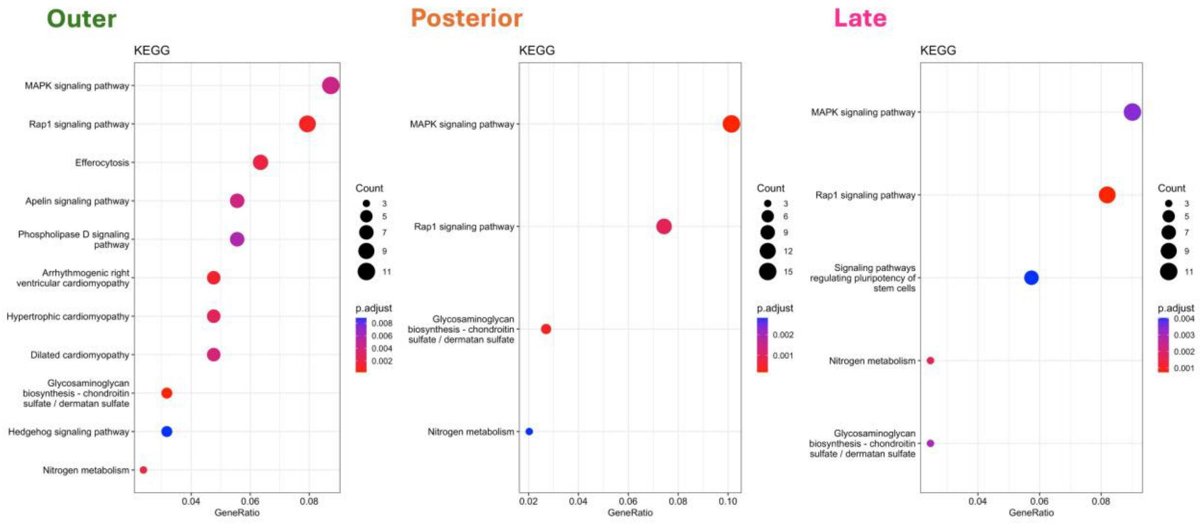


**Supplementary Figure S3 – KEGG pathway analysis reveals MAPK signalling to be more active in the competent region than in any of the non-competent territories.** The transcriptional profiles of the studied regions were compared by KEGG pathway analysis. Here we show the pathways that are enriched in competent epiblast versus each of the non-competent regions (Outer, Posterior and Late). Note that in each case the top hit represents components or targets of MAPK signalling, suggesting that this pathway is more active in the competent area opaca than in any of the other regions.

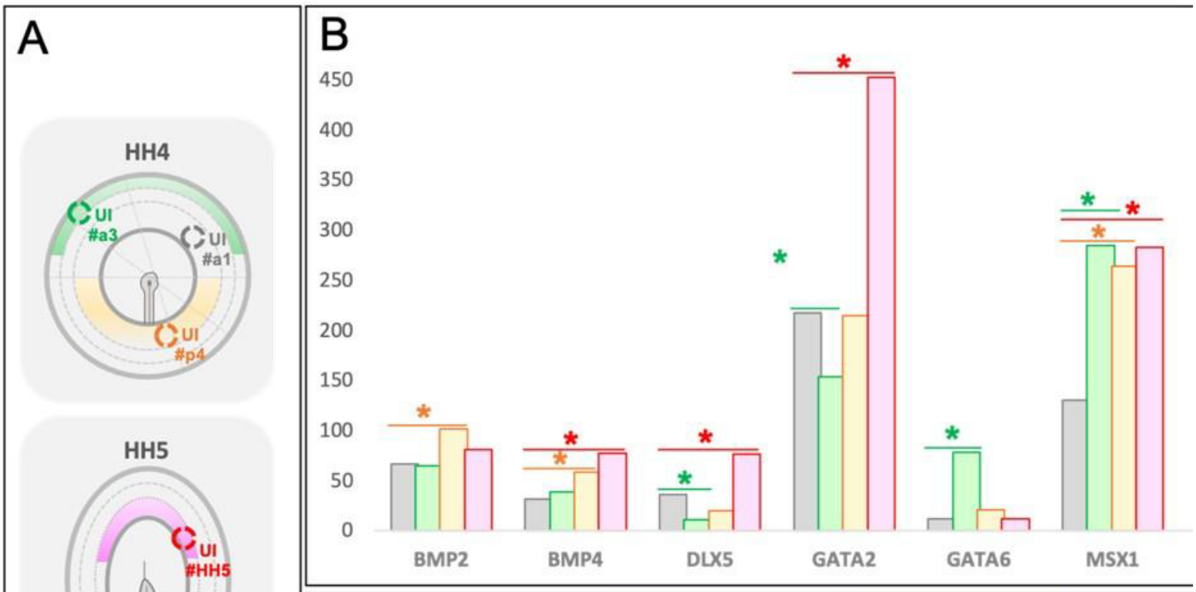


C

| RNA sq | a1 mean | a3 mean | a3 T.Test | p4 mean | p4 T.Test | HH5 mean |
|---|---|---|---|---|---|---|
| BMP2 | 66.4 | 64.5 | 1 | 101.2 | 0.03 | 81.1 |
| BMP4 | 31.6 | 38.7 | 0.5 | 58.7 | 0.01 | 77.3 |
| DLX5 | 36.4 | 11.3 | 0.08 | 19.9 | 0.2 | 76.8 |
| GATA2 | 217.2 | 153.5 | 0.03 | 215.1 | 1 | 452.4 |
| GATA6 | 12.3 | 78.6 | 0.001 | 21.3 | 0.2 | 11.5 |
| MSX1 | 130.1 | 285.1 | 0.07 | 264.5 | 0.03 | 282.9 |

**Supplementary Figure S4 – RNAseq analysis of BMP-related genes. A.** Competent epiblast #a1 (grey) and three non-competent regions: 'outer' (green), 'posterior' #p4 (orange) and 'late' #HH5 (pink) that had not been exposed to an organizer graft ("uninduced", UI) were dissected and subjected to bulk mRNA sequencing. **B-C.** Expression of BMP ligands *BMP2* and *BMP4*, and BMP targets *DLX5, GATA2, GATA6, MSX1* and *SMAD6* was assessed by RNAseq in competent and non-competent regions of area opaca epiblast. RNA counts are plotted as a bar graph (B). The table shows the RNA mean count (fpkm) of the three replicates of each epiblast region and Student's T-Test was calculated comparing all three replicates of each noncompetent epiblast region to the competent epiblast (C). This uncovered statistically significant differences between each non-competent epiblast region and the competent epiblast. Significant differences (p value <0.05) are highlighted in colour in the table (C) and by an asterisk in (B).

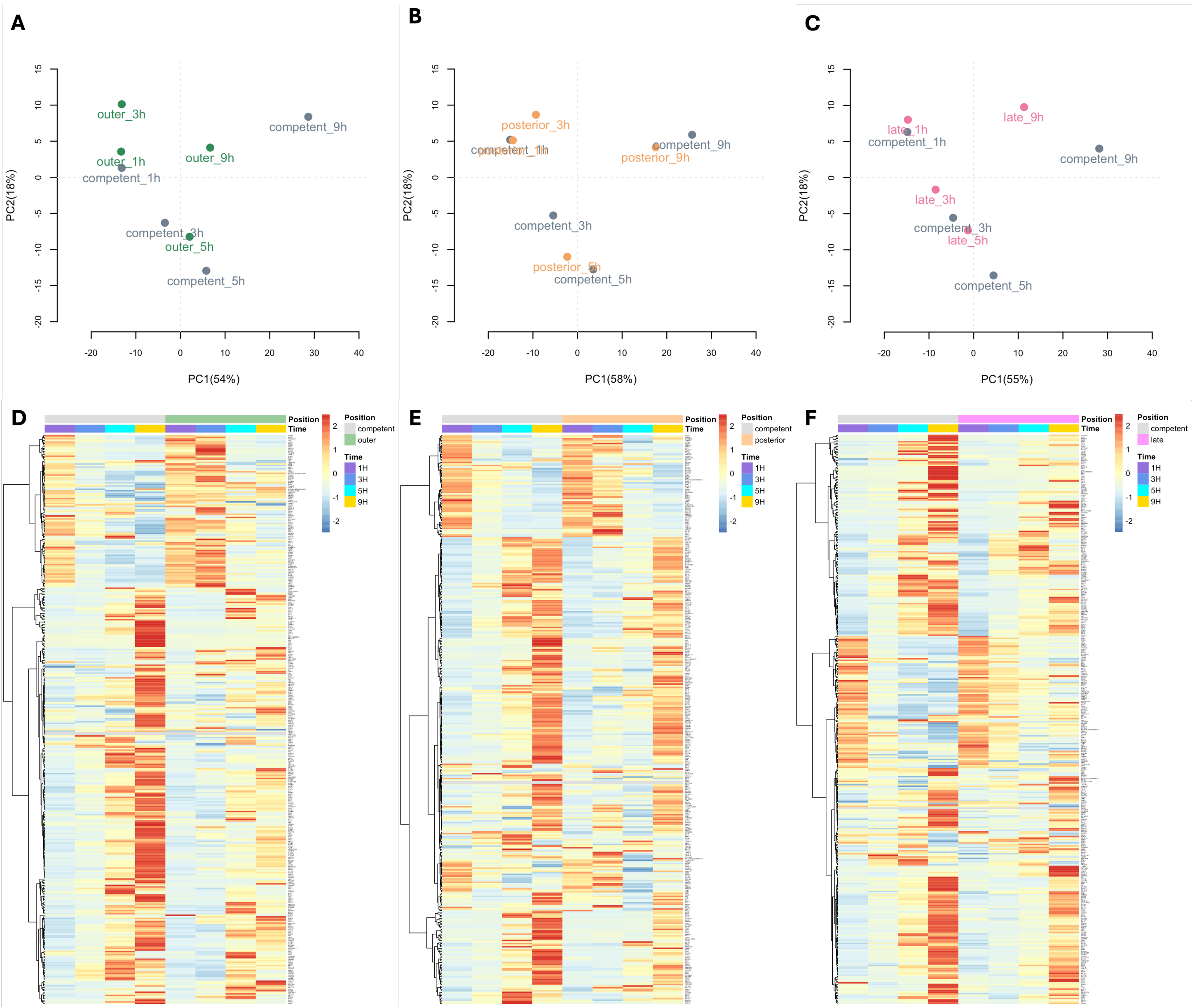


**Supplementary Figure S5 – Time-course analysis of gene expression in the non competent populations** The response to an organizer graft of all the studied regions was analysed in time-course (1, 3, 5 or 9h after the graft) by NanoString nCounter technology using a custom probe set. **A-C.** PCA plots comparing the responses of competent tissue (gray data points) to the outer (green) (A), posterior (orange) (B) and late (pink) (C) at each time point. **D-F.** Differential gene expression of all genes in the custom probe set at every time point for the outer (D), posterior (E) and late (F) regions relative to competent epiblast.

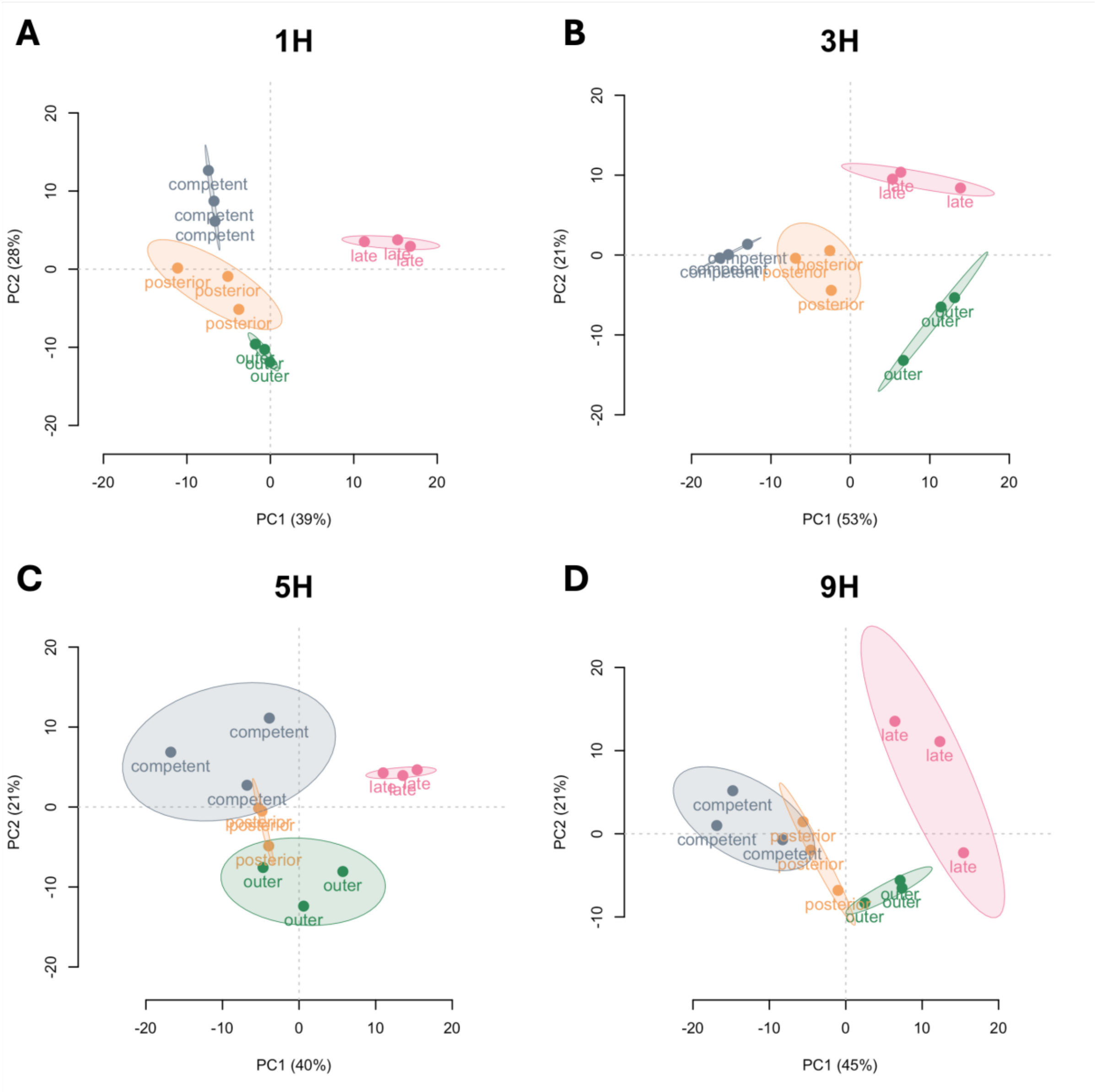


**Supplementary Figure S6 - PCA comparison of the responses to an organizer of the studied regions in time-course.** The responses to an organizer of the competent (grey), outer (green), posterior (orange) and late (pink) regions were analysed in time course identifying gene expression changes after 1h (A), 3h (B), 5h (C) or 9h (D) in the NanoString probe set.

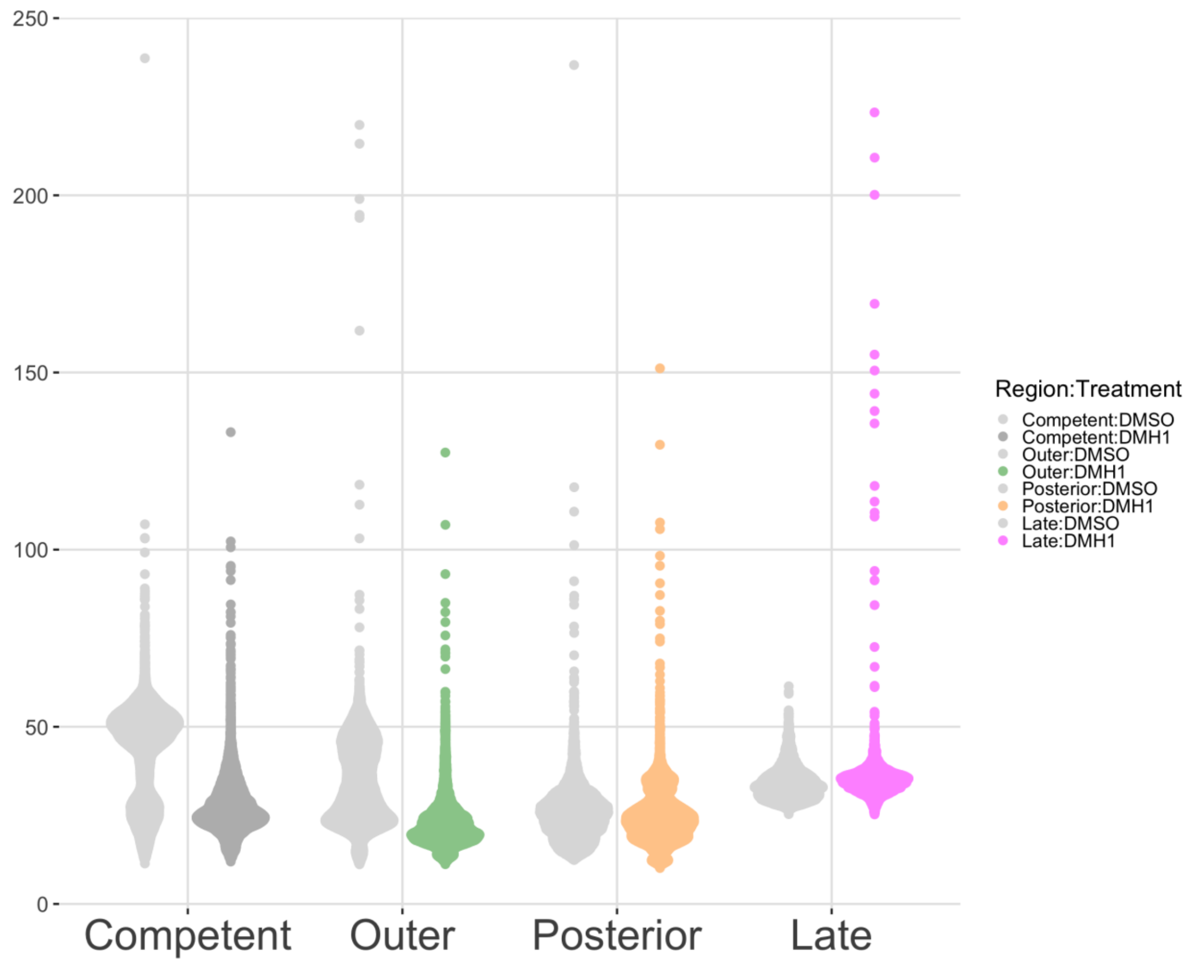


**Supplementary Figure S7 – Quantification of the effects of DMH1.**
Immunostaining of pSmad1/5/8 was performed after 15h incubation in either 2µM DMH1 (the second, coloured, plot for each region) or 0.02% DMSO controls (the first, pale grey plot for each region) on each of the four regions analysed. DMH1 reduces the level of pSmad1/5/8 in the competent, outer and posterior regions relative to DMSO controls.

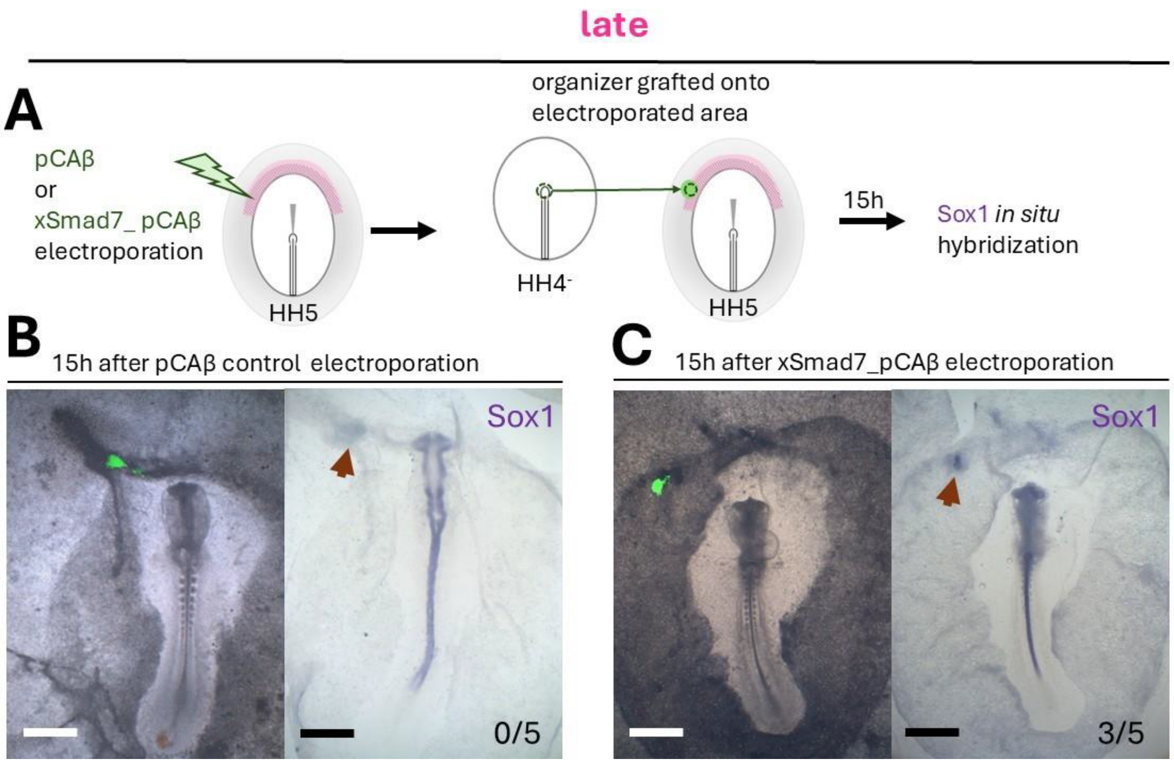


**Supplementary Figure S8 – Inhibition of BMP signalling by Smad7 restores neural competence in the late area opaca.** (A) Schematic representation of the experimental setup. HH5 host embryos were electroporated in the 'late' region of the area opaca with either the control pCAβ vector or xSmad7-pCAβ. An organizer was then grafted onto the electroporated territory, and embryos were incubated for 15 h before fixation and in situ hybridization for *Sox1*. (B, C) Representative embryos showing the electroporated territory (green fluorescence, left) and *Sox1* expression (right). No ectopic *Sox1* expression was detected following control pCAβ electroporation (B; 0/5), whereas *Sox1* was induced following xSmad7-pCAβ electroporation (C; 3/5). Brown arrowheads indicate the electroporated region adjacent to the organizer graft. Scale bars, 500 μm.

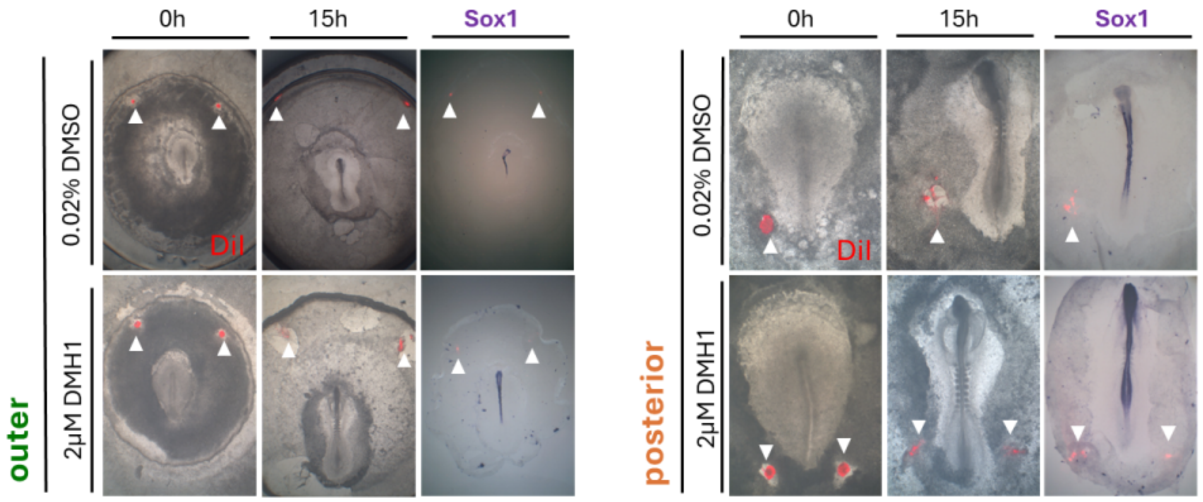


**Supplementary Figure S9 – BMP inhibition by DMH1 is not sufficient to confer competence for neural induction to the outer or posterior regions.** Embryos at stage HH3$^{+}$-4$^{-}$ were incubated for 15h with either 2µM DMH1 or 0.02% DMSO. Organizer grafts (marked with DiI) were placed onto the outer or posterior regions at the beginning of the experiment and expression of the mature neural marker *Sox1* analysed by in situ hybridization. No ectopic *Sox1* expression was seen near the graft site either in outer (0/5 for DMSO controls; 0/5 for DMH1 treatment) or posterior area opaca (0/5 for DMSO controls; 0/5 for DMH1 treatment). Graft sites indicated by arrowheads.

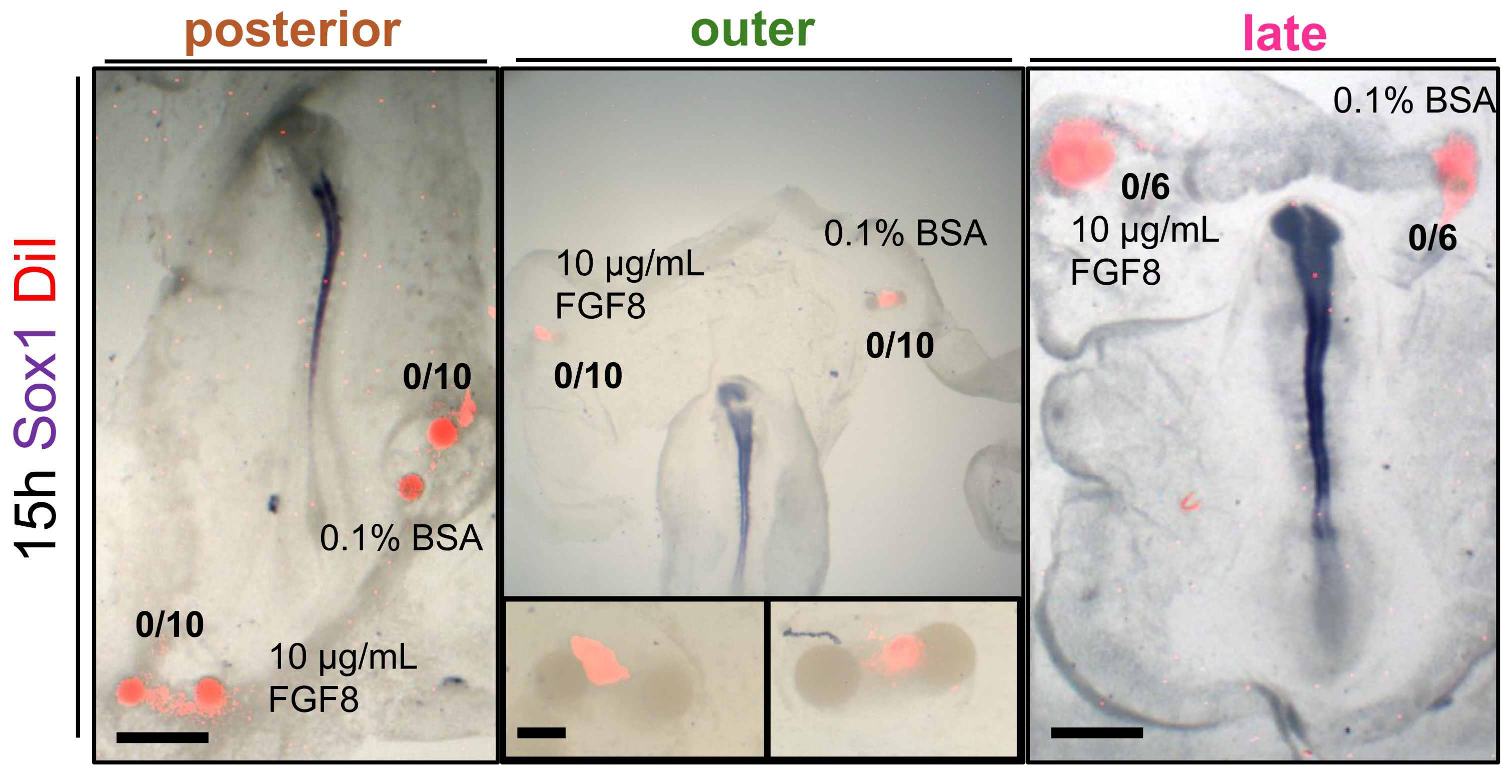


**Supplementary Figure S10 - FGF8 stimulation is not sufficient to induce *Sox1* expression in non-competent territories.** FGF8-coated beads (10 µg/mL) or control beads treated with 0.1% BSA were placed adjacent to organizer grafts in the posterior, outer or late non-competent regions of the area opaca. Organizer grafts were labelled with DiI. Embryos were cultured for 15 h and analysed by in situ hybridization for *Sox1*. No ectopic *Sox1* expression was detected in response to FGF8 treatment in the posterior (0/5), outer (0/5) or late (0/6) regions; control BSA-treated beads likewise failed to induce *Sox1* expression in the corresponding conditions. Scale bars, 500 µm for whole-mount images and 50 µm for insets.

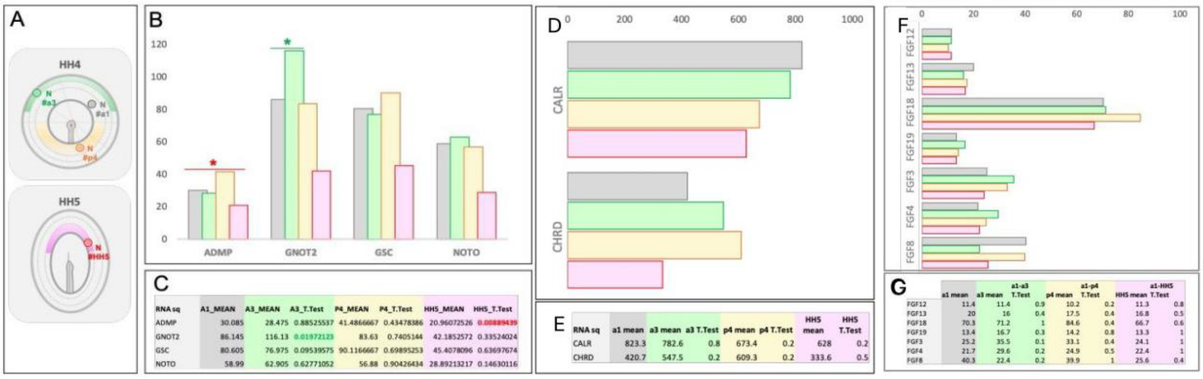


**Supplementary Figure S11 – Transcriptional profile of the organizer after being exposed to non-competent regions.** Five hours after having been grafted onto the competent, outer, posterior or late regions (A), the grafted organizer was collected and studied by RNAseq. Comparison of the expression of node markers (B, p-values of Student's t-test in C), BMP inhibitors (D, with p-values of Student's t-test in E) or FGF signalling molecules (F, p-values of Student's t-test in G) did not reveal significant changes, with the exception of a reduction of *ADMP* after exposure to the late region and an increase of *GNOT2* after grafting onto the outer region (B and C).

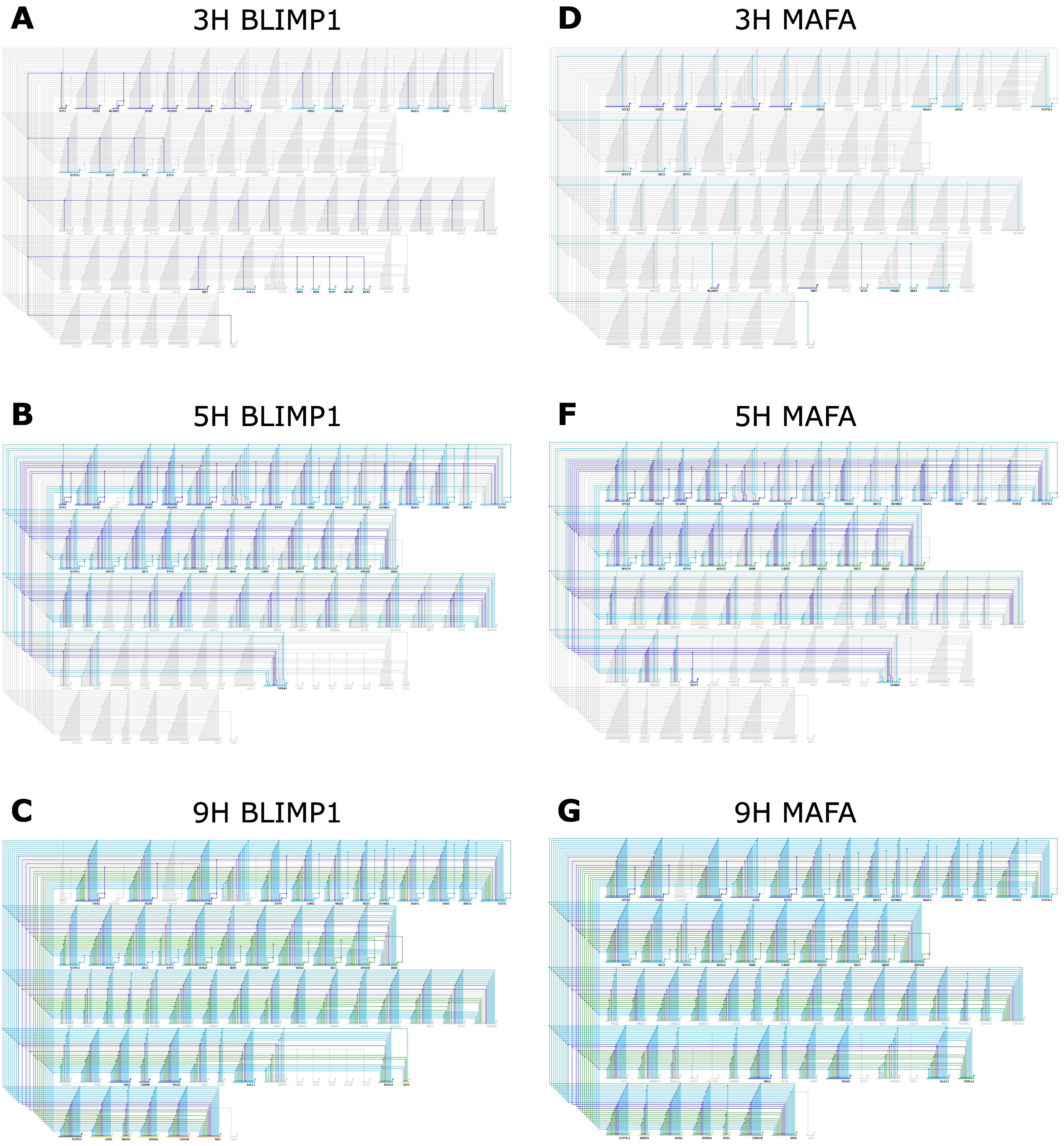


**Supplementary Figure S12 – Targets of *BLIMP1* and *MAFA* in the gene regulatory network of neural induction.** The downstream targets of *BLIMP1* (A, B, C) and *MAFA* (D, E, F) at 3, 5 and 9 H were extracted from the neural induction GRN.
